# Modelling eDNA transport in river networks reveals highly resolved spatio-temporal patterns of freshwater biodiversity

**DOI:** 10.1101/2022.01.25.475970

**Authors:** Luca Carraro, Rosetta C. Blackman, Florian Altermatt

## Abstract

1. The ever-increasing threats to riverine biodiversity call for the development of novel approaches for a complete assessment of biodiversity across highly resolved spatial, temporal and taxonomic scales. Past studies on riverine biodiversity patterns were often restricted to spatially scattered data, focused on specific taxonomic groups, and disregarded the temporal dimension, preventing a universal understanding of relationships between biodiversity and stream size across spatial, temporal and taxonomic scales. Recent advances in the joint use of environmental DNA (eDNA) data and novel mechanistic models for eDNA transport in river networks have the potential to uncover the full structure of riverine biodiversity at an unprecedented spatial resolution, hence providing fundamental insights into ecosystem processes and offering a basis for targeted conservation measures.
2. Here, we applied a mechanistic model (i.e., the eDITH model) to a metabarcoding dataset covering three taxonomic groups (fish, invertebrates and bacteria) and three seasons (spring, summer and autumn) for a 740-km^2^ Swiss catchment, sampled for eDNA at 73 sites.
3. Using the mechanistic model, we upscaled eDNA-based biodiversity predictions to more than 1900 individual reaches, allowing an assessment of patterns of *α*- and *β*-diversity across seasons and taxonomic groups at a space-filling, fine scale over the whole network.
4. We found that both predicted *α*- and *β*-diversity varied considerably depending on both season and taxonomic group. Predicted fish *α*-diversity increased in the downstream direction at all seasons, while invertebrate and bacteria *α*-diversity either decreased downstream or was not significantly related to position within network, depending on the season. Spatial *β*-diversity was mostly found to be decreasing in the downstream direction, and this was the case for all seasons for bacteria. Temporal *β*-diversity was mostly found to be increasing downstream. In general, genus richness values predicted by the model were found to be higher than those obtained by directly analyzing the eDNA data. Overall, stream size (subsumed by drainage area) was generally a poor predictor of patterns of predicted *α*- and *β*-diversities. Conversely, riverine biodiversity is shaped by a complex interplay of environmental variables, abiotic and biotic factors, which need be taken into account for a correct assessment of its structure.

## 1 Introduction

Freshwater ecosystems are among the most biodiverse ecosystems worldwide, in relation to their area [Dudgeon, 2020; Vörösmarty et al., 2010], but also among the most threatened with respect to loss of biodiversity [Darwall et al., 2018; Reid et al., 2019]. Strategies for conservation of biodiversity should be based on complete biodiversity assessments across spatial and temporal scales, as well as taxonomic groups in order to fully understand and preserve ecosystem functioning [Altermatt et al., 2020]. However, this is often not the case for river systems, due to the spatial structure of riverine metacommunities, the coarse spatio-temporal resolution of biodiversity data, a limited taxonomic coverage, and the difficulty to transfer knowledge from one taxonomic group to another [Barbour, 1999; Altermatt, 2013; Altermatt et al., 2020; Darwall et al., 2011].

A seminal model for ecological communities in rivers, the river continuum concept [Vannote et al., 1980], predicted species diversity to have a unimodal patterns in very large rivers (up to 12^th^ Strahler order), with the highest richness observed in mid-order reaches. For most rivers of intermediate size, this translates into an increasing pattern of *α*-diversity in the downstream direction, which has been validated empirically [Ward, 1998], in particular for fish [Muneepeerakul et al., 2008] and macroinvertebrates [Altermatt et al., 2013; Tonkin et al., 2015; Blackman et al., 2021b]. Conversely, bacteria richness was generally found to follow a decreasing trend in the downstream direction [Besemer et al., 2013; Ruiz-González et al., 2015; Savio et al., 2015]. The other component of total (*γ*-) diversity, namely *β*-diversity, has been less often investigated, although Finn et al. [2011] observed decreasing *β*-diversity with increasing stream size in macroinvertebrates. However, universal relationships between biodiversity and stream size appear to be elusive [Vander Vorste et al., 2017]. Crucially, most of the studies investigating biodiversity patterns in rivers only focused on specific taxonomic groups, or neglected the temporal dimension of biodiversity, either by considering a snapshot of data collected at a single time point, or by analyzing temporally averaged data. Moreover, most biodiversity studies have been based on spatially scattered, pointwise data, hence preventing a spatially highly resolved and/or space filling assessment of biodiversity.

In this perspective, environmental DNA (eDNA, i.e. DNA isolated from environmental samples [Taberlet et al., 2012; Pawlowski et al., 2020]) has opened new avenues for fast, cost-effective and taxonomically broad biodiversity assessments [Thomsen and Willerslev, 2015; Valentini et al., 2016; Deiner et al., 2017; Beng and Corlett, 2020]. In particular, eDNA increases our understanding of biodiversity structure and related ecosystem processes in riverine ecosystems [Altermatt et al., 2020], especially considering that, due to downstream transportation of DNA molecules with streamflow, eDNA constitutes an aggregated measure of biodiversity across large drainage areas [Deiner and Altermatt, 2014; Barnes and Turner, 2015; Deiner et al., 2016; Shogren et al., 2017; Seymour et al., 2021]. Correct interpretation of eDNA data collected in rivers hence requires consideration of the role of hydrological transport and decay of genetic material. The recently developed eDITH model (*eD*NA *I*ntegrating *T*ransport and *H*ydrology) couples a geomorphological and hydrological characterization of a catchment, eDNA transport and decay dynamics, and a species distribution model, and allows transforming pointwise eDNA data collected at a catchment into predicted maps of taxon density [Carraro et al., 2017, 2018, 2020b, 2021]. The eDITH model has hitherto been successfully applied to predict the distribution of single species, i.e. a fish parasite and its primary host [Carraro et al., 2017, 2018], as well as biodiversity of aquatic insects at a given point in time [Carraro et al., 2020b]. Given its generality and the basic assumptions on eDNA shedding and decay processes underpinning its formulation [Carraro et al., 2018], the eDITH model can in principle be applied to any taxonomic group, and can also be used to identify temporal variations in biodiversity patterns in river networks.

Here, we applied the eDITH model to an eDNA metabarcoding dataset covering three taxonomic groups relevant to freshwater communities (fish, invertebrates and bacteria), and three seasons (spring, summer and autumn), providing predictions of patterns of *α*- and *β*-diversity, and changes thereof with respect to season and taxonomic groups, at an unprecedented space-filling, highly resolved spatial scale (∼500-m long river reaches) covering a 740-km^2^ Swiss catchment [Blackman et al., 2021a].

## 2 Methods

### 2.1 Study area

The Thur (Fig. 1) is a pre-alpine river located in northeastern Switzerland, draining an area of 740 km^2^, for which extensive data on hydrology and biodiversity are available [Abbaspour et al., 2007; Mächler et al., 2019; Carraro et al., 2020b; Blackman et al., 2021a]. Here, eDNA samples were collected at 73 sites in three different seasons in 2018: spring (17^th^-24^th^ May), summer (20^th^ Aug-5^th^ Sep), and autumn (2^nd^-8^th^ Oct). In summer, 4 sites were not sampled because their respective reaches were temporarily dry. Hydrological data on the catchment were available at four gauging stations (Fig. 1) operated by the Swiss Federal Office for the Environment, while landscape data on elevation, land cover types and geology were provided by the Swiss Federal Office for Topography (see Carraro et al. [2020b] for details).

**Figure 1:**
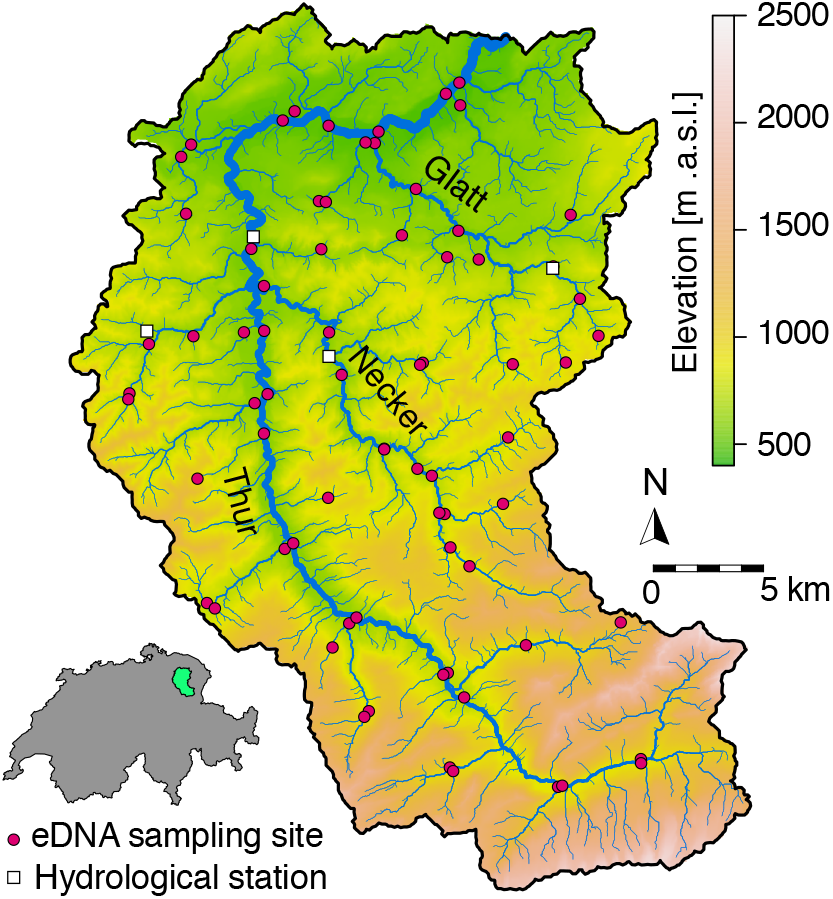
Map of the Thur catchment displaying the locations of eDNA sampling sites (red dots) and hydrological stations (white squares). The three main tributaries (rivers Thur, Glatt and Necker) are also identified.

The river network was extracted by using a TauDEM implementation of the D8 method [O’Callaghan and Mark, 1984] on a 25-m digital elevation model of the region. A threshold drainage area of 0.25 km^2^ was used to identify the sources of the river network, resulting in a total river length of 1034 km. Such threshold area value was chosen as the maximum one such that all 73 sampling sites effectively belonged to the resulting river network. The river was partitioned into reaches of maximum length 1 km, by following Carraro et al. [2020a], which resulted in a total of 1908 reaches, with mean length 542 m.

### 2.2 eDNA data collection and sequencing

Details on eDNA data collection and sequencing are elucidated in Blackman et al. [2021a], but are here briefly recapitulated in order for this work to be self contained.

Environmental DNA samples were collected and filtered on site using disposable 50 mL syringes and 0.22 *μ*m sterivex filters (Merck Millipore, Merck KgaA, Darmstadt, Germany). At each site 1 L of water was filtered. Samples were extracted in clean lab facilities using the DNeasy PowerWater Sterivex Kit (Qiagen, Hilden, Germany) following the manufacturer’s protocol. Three libraries were constructed for the following markers: 12S, COI and 16S using a two-step (12S and COI) and three-step (16S) library preparation method, where clean amplicons were indexed using unique combinations of the Illumina Nextera XT Index Kit A, C and D in the last PCR, following the manufacturer’s protocol (Illumina, Inc., San Diego, CA, USA). Paired-end sequencing was performed on an Illumina MiSeq (Illumina, Inc. San Diego, CA, USA) at the Genetic Diversity Centre at ETH, Zurich (see Blackman et al. [2021a] for full details of each library preparation). After each of the libraries were sequenced, the data was demultiplexed and reads were quality checked. Raw reads were end-trimmed, merged and quality filtered, additional reads were clustered at 99% identity to obtain error corrected and chimera-filtered sequence variants ZOTUs. The final ZOTUs were then clustered using a 97% similarity approach and taxonomic assignment with a 0.85 confidence threshold. Taxonomic assignment was carried out with the following databases: 12S: NCBI BLAST (v200416), COI: Custom reference database (Including MIDORI un-trimmed (V20180221) and 16S: SILVA (V128). Prior to data analysis a 0.1% contamination threshold was applied to each sample, species with a non-aquatic life stage were removed from the data set and the data was merged at genus level.

Overall, we detected 12 fish, 80 invertebrate and 282 bacterial genera. The three fish genera *Barbus, Gobio* and *Phoxinus* were detected with both the 12S and COI barcode regions. To avoid duplicated coverage, we removed from the database the read numbers corresponding to these three genera from the COI library, as the corresponding read numbers in the 12S library were higher and the latter marker region was used to target fish. Fig. S1 displays the total number of reads observed and the total number of genera detected for each barcode region, taxonomic group and season, pooled over all sampling sites.

### 2.3 eDITH model

The eDITH model implementation essentially follows Carraro et al. [2020b], and is here summarized for the specific study setting. For each genus the expected eDNA concentration *C*_*j*_ at a sampling site *j* of the network reads:

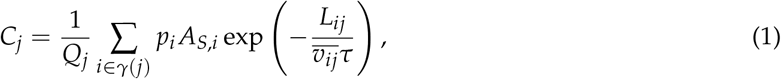

where *Q*_*j*_ is the water discharge at reach *j* (i.e., the reach where sampling site *j* is located), *γ*(*j*) identifies the set of reaches upstream of *j* (with *j* included), *p*_*i*_ the eDNA production rate at reach *i, A*_*S,i*_ the source area of reach *i* (namely its open water surface), *L*_*ij*_ the along-stream path from *i* to 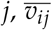 the average water velocity along such path, *τ* a characteristic decay time. eDNA production rates are expressed via a Poisson generalized linear model as *p*_*i*_ = *p*_0_ exp (***β***^*T*^**X**(*i*)), where **X**(*i*) is a vector of environmental covariates, ***β*** a vector of covariate effects and *p*_0_ a baseline production rate. We utilized 35 covariates, representing morphological, land cover, geological and geographical characteristics of the catchment. These covariates correspond exactly to those used in Carraro et al. [2020b]. Observed read data from each genus at a given site *j* and a given season were assumed to follow a geometric distribution, with mean proportional to *C*_*j*_. Following Carraro et al. [2020b], reach width was evaluated via aerial images in correspondence to the four hydrological stations, and a power-law relationship with drainage area was derived. Width values were then extrapolated to all 1908 reaches via the so-obtained power-law relationship. The same procedure was performed for the three different seasons for discharge and water depth, whose data values were taken as the averages of the mean daily measured values at the hydrological stations during the respective sampling periods. A power-law relationship on drainage area was then fitted separately for each hydrological variable (i.e., discharge, water depth) and season, and then extrapolated to the whole catchment. Finally, we calculated water velocity values at all reaches for all seasons under the hypothesis of rectangular river cross-sections (i.e., *v* = *Q*/(*wd*), where *v* is velocity, *Q* is discharge, *w* is width and *d* is depth).

The posterior distributions of the 37 unknown parameters (i.e., vector ***β*** containing effect sizes for 35 covariates, decay time *τ* and baseline production rate *p*_0_) were inferred independently for each season and genus, by using the DREAM_*ZS*_ [Vrugt et al., 2009] *algorithm, implemented via the BayesianTools* R-package [Hartig et al., 2019]. Three independent Markov chains were run, with a total chain length of 3 · 10^6^ (plus a burn-in length of 5 · 10^5^). A normal prior distribution with null mean and standard deviation of 3 was adopted for all ***β*** components; *p*_0_ had a uniform prior bound between 0 and 1; a log-normal prior for *τ* was chosen, with a median of 5 h and a mode of 4 h. The so-obtained maximum a posteriori parameter estimates were used to produce maps of relative species density (i.e., *p*_*i*_). These were subsequently translated into detection probability maps by evaluating the expected read number that would be observed at a reach if the reach were disconnected from the river network, and by assessing the probability that the measured read number therein would be larger than 0 according to the assumption of geometric distribution of read numbers (see Carraro et al. [2020b] for details). Finally, presence/absence maps for each genus were derived by imposing a threshold of 0.5 on detection probability.

### 2.4 Evaluation of *α*- and *β*-diversity patterns

For each taxonomic group and season, the number of genera predicted by the eDITH model to be present in each of the 1908 reaches was taken as a measure of *α*-diversity. We then performed a linear regression to assess the effect of drainage area on genus richness. Given that values of genus richness at the different reaches are in principle not independent (i.e., due to the spatial structure of the covariates used to predict the taxon patterns, predicted *α*-richness values for nearby reaches tend to be correlated), we refrained from performing classic statistical tests on the slope of the linear regression. Instead, we assessed the significance of the effect of drainage area via a bootstrapping approach: we subsampled 500 out of 1908 reaches in a quasi-random fashion (i.e., by splitting reaches into 10 bins according to the drainage area deciles, and randomly sampling (with replacement) 50 reaches within each bin), and linearly regressed genus richness on drainage area for the subsampled reaches. We repeated this procedure 100 times, and considered a positive (respectively negative) significant effect of drainage area if, in at least 95 out of 100 cases, the fitted slope of the linear regression was positive (respectively negative). The 2.5^th^-97.5^th^ percentile range of the so-obtained 100 linear regression lines was used as confidence interval of the linear model fit. Moreover, we also computed genus richness from the raw eDNA data at the 73 sampling sites across seasons and taxonomic groups.

In order to evaluate spatial patterns of *β*-diversity with respect to each taxonomic group, the 1908 reaches were partitioned into two location groups, i.e. upstream, if their drainage area was lower than the median value across all reaches, or downstream, otherwise. Within each location group, we picked pairs of flow-unconnected sites such that each site appeared in only one pair; the choice of pairs was operated randomly, and was stopped when no other pair could be formed from the sites that had not been picked yet. Note that we chose to limit our attention to *β*-diversity of flow-unconnected sites in order to correct for the fact that downstream sites are more likely to be connected by flow than upstream sites (and hence inherently more prone to show similar community compositions), since the latter mostly consist of headwater reaches. Moreover, picking each site only once ensures that measures of *β*-diversity among all pairs are mutually independent. For each so-obtained pair, Jaccard distance was evaluated via the *betapart* R-package [Baselga and Orme, 2012], which also allowed partitioning of nestedness and turnover components of total *β*-diversity. We accounted for the stochasticity in the choice of pairs by repeating the pair selection process 100 times. The effect of location was deemed significant if the equal-tailed 95% confidence interval of the mean Jaccard distance across one location group did not overlap with that of the other group. Given the relatively limited number (73) of sampling sites available, we found it unfeasible to repeat the aforementioned procedure to evaluate spatial *β*-diversity patterns for the raw eDNA data.

Finally, temporal patterns of *β*-diversity were evaluated by comparing, within each taxonomic group, predicted presence/absence for all taxa at different seasons via the Jaccard distance evaluated at every reach. In particular, we treated the spring season as a benchmark and focused on patterns of spring-to-summer and spring-to-autumn temporal *β*-diversity. We then linearly regressed these patterns against drainage area to possibly detect an upstream/downstream gradient on temporal *β*-diversity. In order to assess the significance of such trends, we adopted the same bootstrapping procedure that was earlier described with respect to *α*-diversity. Moreover, we computed temporal *β*-diversity (expressed as Jaccard distance) for the raw eDNA data across seasons and taxonomic groups.

## 3 Results

### 3.1 *α*-diversity patterns

Overall, patterns of *α*-diversity across the different taxonomic groups and seasons show weak, and often non-significant relationships with drainage area (Table 1, Fig. 2). Indeed, the proportion of variance in *α*-diversity explained by drainage area is in all cases lower than 4%, with the exception of fish in spring, where drainage area explains 16.6% of the variance in *α*-diversity (Fig. 2). Conversely, patterns of *α*-diversity appear to be driven by clusters of neighbouring reaches (Fig. 3), which is reflected by the role of environmental covariates in explaining the spatial patterns of taxa (see Fig. S2 for a summary of significant covariates depending on taxonomic groups and seasons). In general, genus richness values predicted by the eDITH model are higher than those obtained by directly analyzing the eDNA data (Fig. 2, insets) especially for invertebrates (irrespective of the season), and in summer also for fish and bacteria. Moreover, for fish and invertebrates and irrespective of the season, the eDITH model predicts higher *α*-diversity with respect to the raw data for low drainage areas (Fig. 2).

**Table 1:** Summary of the effect of drainage area (i.e. position within network) on *α*- and *β*-diversity patterns. ↗: increasing in the downstream direction; ↘: decreasing in the downstream direction; →: invariant relationship; : significant relationship; ^*NS*^: non-significant relationship. Approaches to detect significance are detailed in the Methods.

| | $\alpha$ diversity | | | Spatial $\beta$ diversity | | | Temporal $\beta$ diversity | |
| --- | --- | --- | --- | --- | --- | --- | --- | --- |
|  | spring | summer | autumn | spring | summer | autumn | spr.-sum. | spr.-aut. |
| Fish | $\nearrow^*$ | $\nearrow^*$ | $\nearrow^*$ | → | $\searrow^*$ | $\nearrow^*$ | $\nearrow^*$ | $\nearrow^*$ |
| Invertebrates | $\searrow^*$ | $\searrow^*$ | $\nearrow^{NS}$ | → | → | $\searrow^*$ | $\nearrow^*$ | $\nearrow^*$ |
| Bacteria | $\nearrow^{NS}$ | $\searrow^*$ | $\nearrow^{NS}$ | $\searrow^*$ | $\searrow^*$ | $\searrow^*$ | $\nearrow^*$ | $\nearrow^{NS}$ |

**Figure 2:**
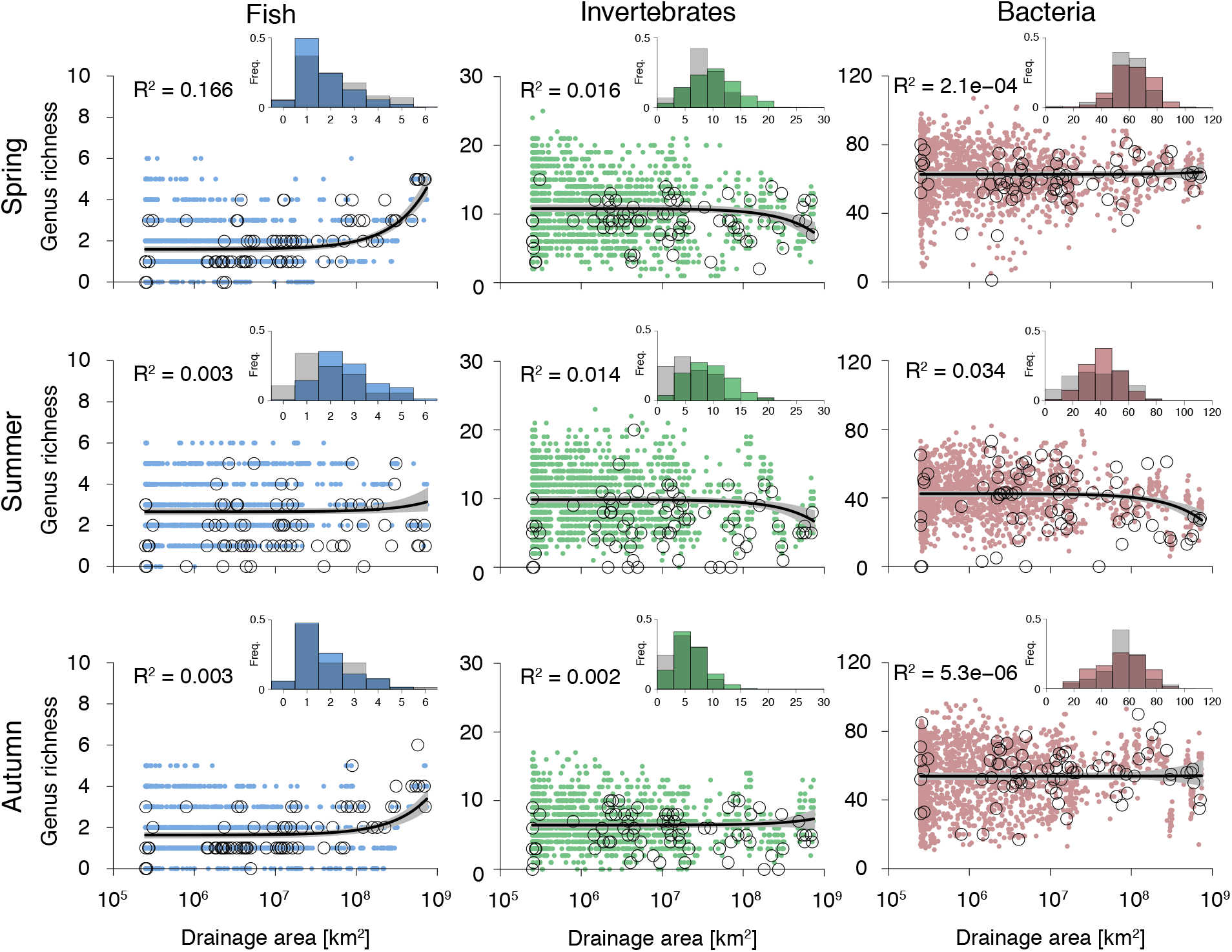
Patterns of *α*-diversity as a function of drainage area. Colored, closed dots represent eDITH model results; for comparison, values inferred from eDNA data are displayed with black, open dots. Black solid lines represent linear model fits on modelled genus richness (*R*^2^ values are reported on the top-left corner). Shaded areas represent 95% confidence intervals on linear model fit (obtained via a bootstrapping technique detailed in the Methods). Note that linear models were fitted on natural values on drainage area, hence the trend lines are exponential in these semi-logarithmic plots. Inset: frequency distribution of genus richness values as predicted by eDITH (colored bars) vs. inferred from eDNA data (grey bars).

**Figure 3:**
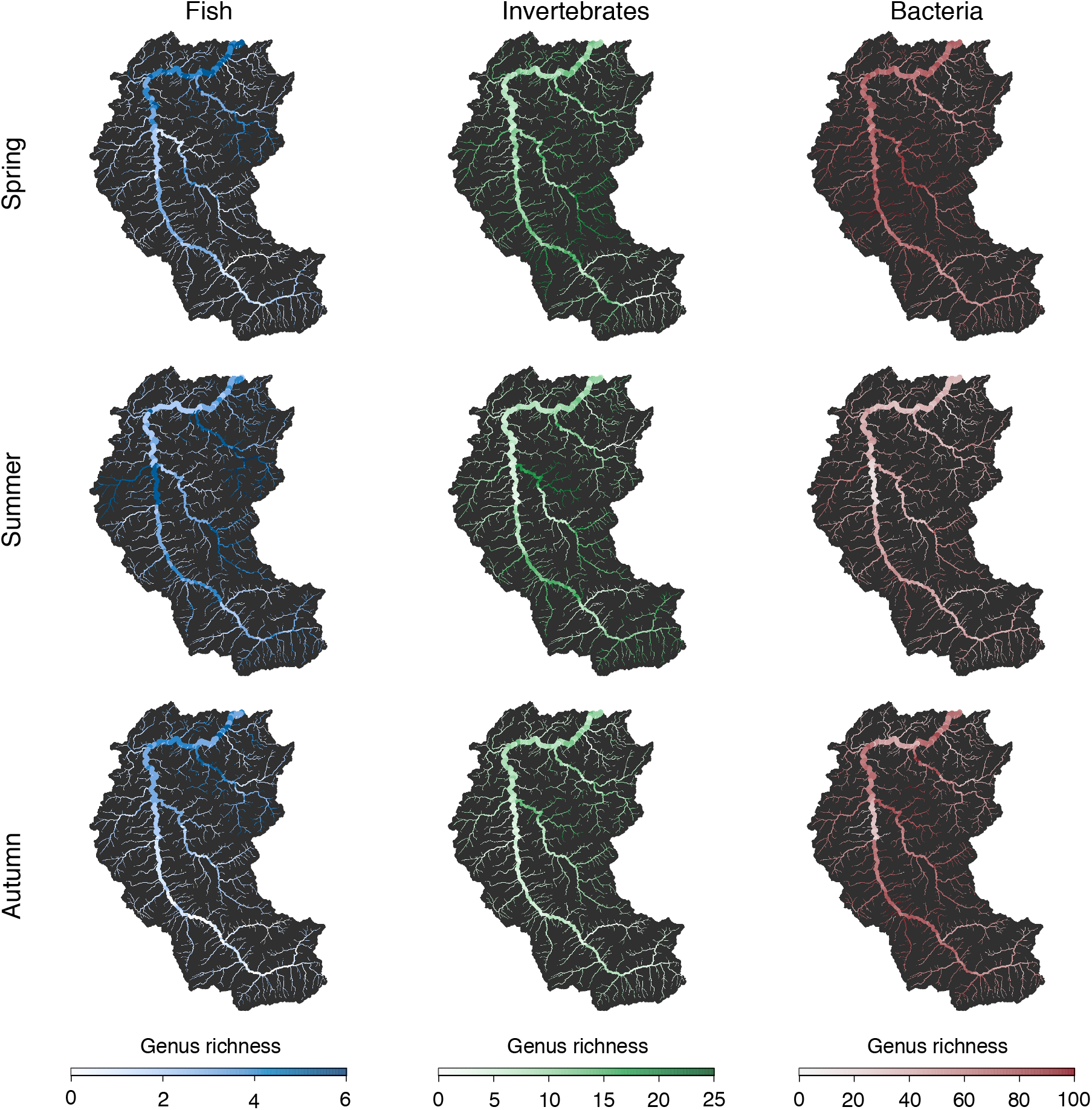
Spatial patterns of predicted *α*-diversity (expressed in terms of genus richness) for the different taxonomic groups and seasons. Displayed values correspond to colored, closed dots in Fig. 2.

Predicted fish *α*-diversity is significantly correlated with drainage area at all seasons (Table 1, Fig. 2). While fish genus richness in spring and autumn is mostly concentrated in the downstream reaches of the watershed, richness in summer is highest at small tributaries at intermediate distance from the river outlet (Fig. 3). For invertebrates, the trend of predicted *α*-diversity decreases significantly in the downstream direction in spring and summer, while it increases non-significantly in autumn (Table 1, Fig. 2). Clusters of high invertebrate *α*-diversity are predicted in the mid-Thur and Necker reaches, which is in qualitative agreement with the patterns found by Carraro et al. [2020b] for the orders Ephemeroptera, Plecoptera and Trichoptera in late June (note that, in the data set here analyzed, 29 invertebrate genera out of 80 belong to one of these three orders). Invertebrate genus richness in autumn tends to be lower as compared to spring and summer (Figs. 2, 3). This finding is reflected by the sensibly lower number of reads observed in autumn for invertebrates as compared to the other seasons, although the total number of invertebrate genera detected in autumn (56) is comparable to that of spring (62) and summer (57) (see Fig. S1). Predicted bacteria *α*-diversity shows a flat distribution across the catchment for all seasons, with a significant decrease in the downstream direction observed only in summer, while the effect of drainage area is not significant in spring and autumn. Summer values of bacterial *α*-diversity are considerably lower with respect to spring and autumn (Figs. 2, 3).

Patterns of predicted *α*-diversity for any taxonomic group are positively correlated across seasons, with spring-autumn correlations being higher than spring-summer correlations for all taxonomic groups (Fig. S3). Moreover, invertebrate *α*-diversity is strongly correlated with bacteria *α*-diversity at all seasons, suggesting a link between these contiguous trophic levels; fish *α*-diversity also shows positive correlations with invertebrate *α*-diversity for all seasons, although the effect is in this case less strong (Fig. S4).

### 3.2 *β*-diversity patterns

The effect of drainage area on predicted spatial *β*-diversity patterns depends greatly on the taxonomic group and the season (Table 1, Fig. 4). For fish, spatial *β*-diversity decreases in the downstream direction in summer, while the pattern is reversed in autumn, and no effect of drainage area is observed in spring; for invertebrates, there is no significant trend of drainage area on spatial *β*-diversity in spring and summer, while a decreasing trend is observed in autumn; finally, spatial *β*-diversity of bacteria decreases significantly with drainage area for all seasons. Importantly, across all seasons, values of the Jaccard distance for invertebrates are much larger than those for fish and bacteria. Analysis of the relative contribution of nestedness and turnover components to the total Jaccard distance shows the predominant role of turnover in *β*-diversity for invertebrates and bacteria across all seasons, while a larger role of nestedness is observed for fish (Table S1).

**Figure 4:**
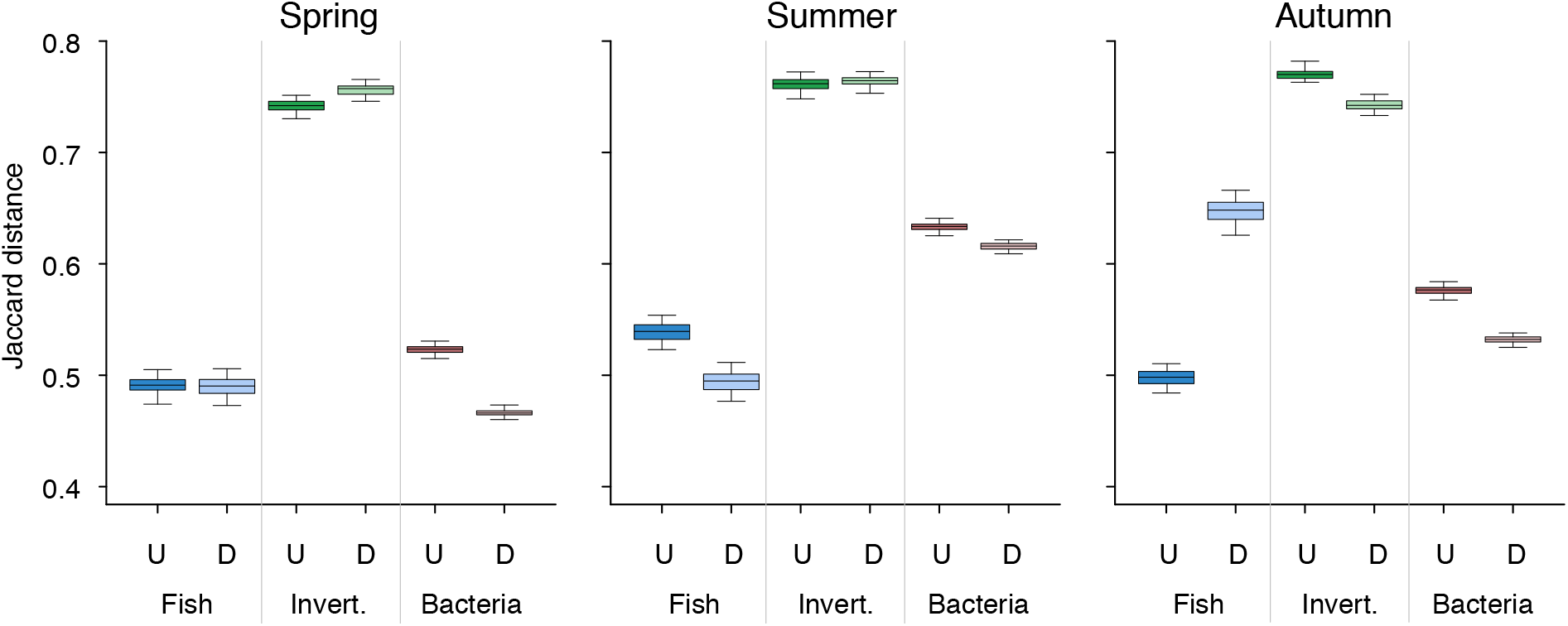
Effect of drainage area on predicted spatial *β*-diversity of fish, invertebrates (“Invert.”) and bacteria, respectively. Each boxplot contains 100 values, representing the mean pairwise Jaccard distance for one of the 100 replicated choices of pairs within a group of reaches (“U”: upstream; “D”: downstream - see Methods). Boxes’ extent corresponds to interquantile range; whiskers’ extent to 2.5^th^-97.5^th^ percentile range.

Patterns of predicted temporal *β*-diversity are significantly and positively related to drainage area for all taxonomic groups and seasons, with the exception of bacteria in the spring-autumn comparison, where the postive trend is not significant (Table 1, Fig. 5). However, the proportion of variance in Jaccard distance explained by drainage area is low (i.e., consistently lower than 8%). Interestingly, for all taxonomic groups, higher values of predicted temporal *β*-diversity are observed in the spring-summer, than in the spring-autumn comparison (Table S2). This reflects the fact that patterns of spring *α*-diversity are better correlated to patterns of autumn, than summer *α*-diversity (Fig. S3). Moreover, values of temporal *β*-diversity for invertebrates are higher than those for fish and bacteria, thus showing that predicted patterns of invertebrate communities (or their detectability, see Discussion) are much more diverse in both spatial and temporal dimensions than other taxonomic groups. For fish, larger values of Jaccard distance in the spring-summer comparison are observed at the downstream reaches, while the highest temporal *β*-diversity is found at the mid-to-upper Thur reaches (Fig. 6). Regarding invertebrates, the predicted spatial trends of temporal *β*-diversity tend to be more constant across seasons, with larger values observed along the main Thur stem, and more stable communities in the Necker and Glatt tributaries (Fig. 6). Bacteria temporal *β*-diversity shows a less pronounced spatial variability, with all values being larger than 0.4 at all seasons (Fig. 5). As it was the case for spatial *β*-diversity, temporal *β*-diversity of fish is mostly driven by nestedness, while turnover plays a major role in the temporal *β*-diversity of invertebrates and bacteria (Table S2). Finally, bacteria temporal *β*-diversity evaluated on the raw eDNA data is considerably lower with respect to that evaluated via the eDITH model (Fig. 5, insets). As for fish and invertebrates, differences between the two approaches are less marked, although data-based temporal *β*-diversity tends to assume more extreme (both low and high) values.

**Figure 5:**
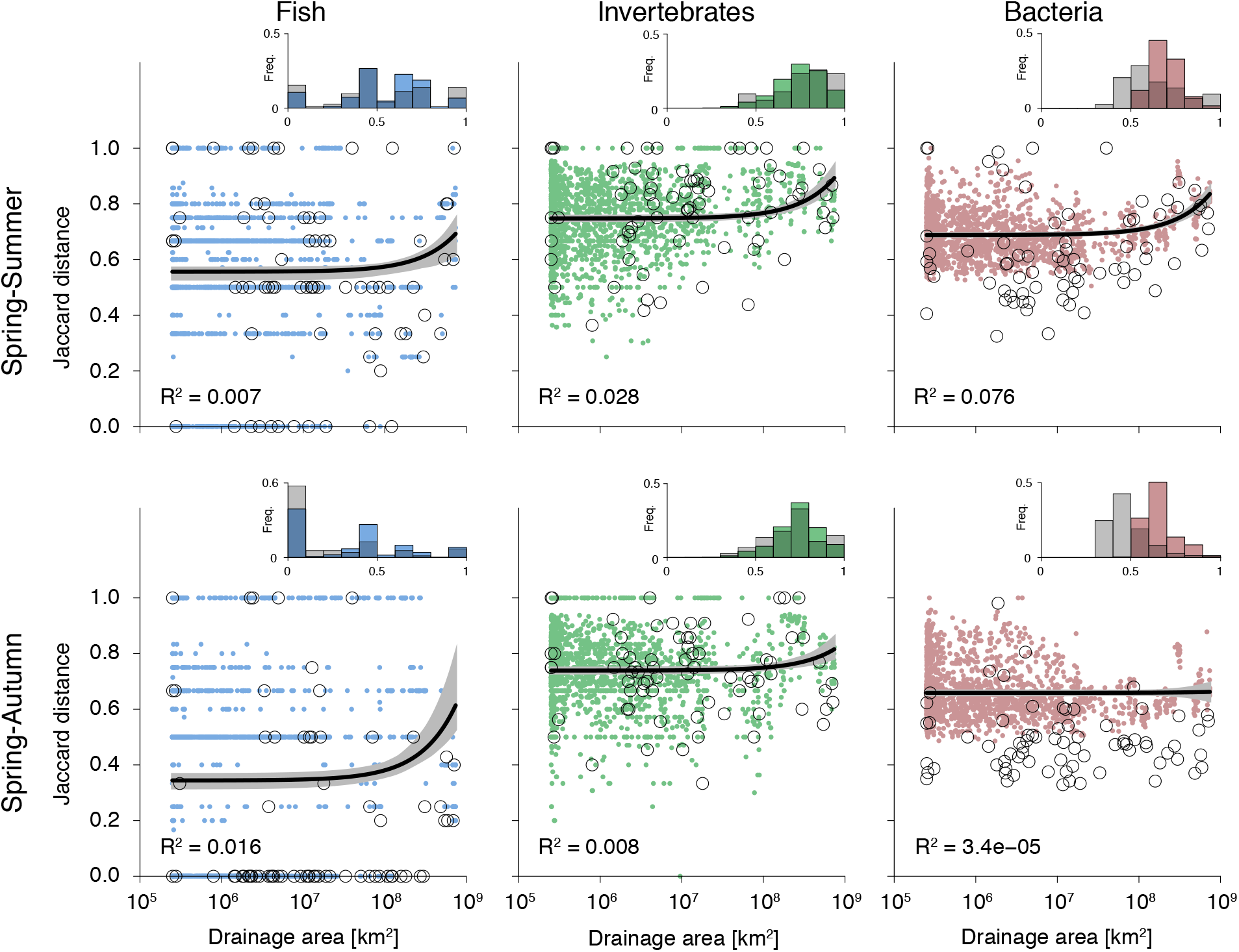
Patterns of predicted temporal *β*-diversity as a function of drainage area. Figure construction is as in Fig. 2.

**Figure 6:**
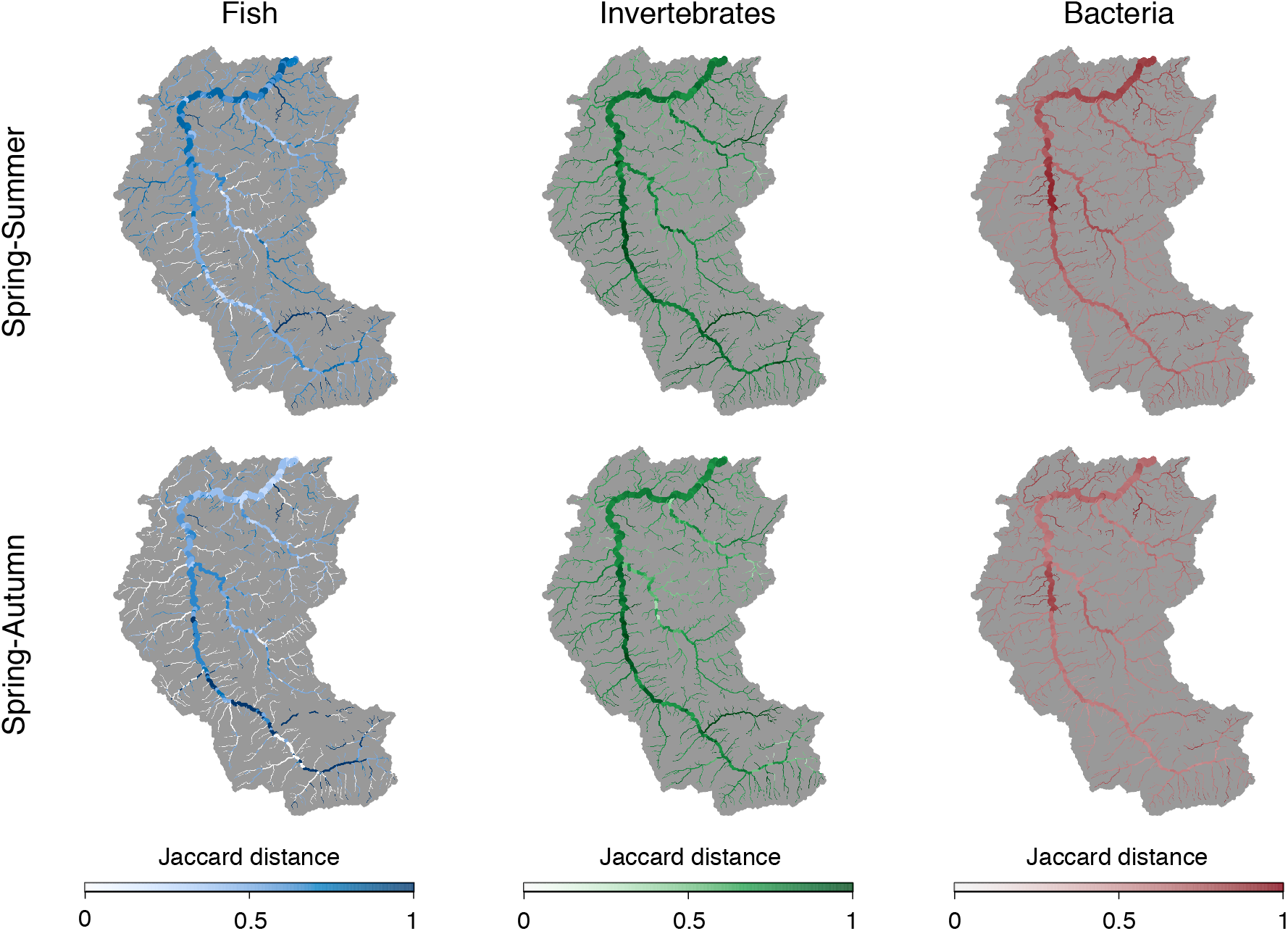
Spatial patterns of predicted temporal *β*-diversity (expressed via the Jaccard distance).

## 4 Discussion

The combined use of the eDITH model and a spatially and temporally replicated multimarker eDNA data set allowed the assessment of patterns of *α*- and *β*-diversity across a river catchment with approximately a 30-fold increased resolution compared to the initial sampling sites. Specifically, using data from 73 sampling sites across the whole catchment, diversity patterns across all taxonomic groups were extrapolated at a genus level to cover a total of 1908 river reaches (of mean length ∼500 m) along the complete river network, giving unprecedented spatially covered information on fish, invertebrates and bacteria. Comparison of these *α*- and temporal *β*-diversity patterns obtained via the eDITH model vs. the raw eDNA data revealed substantial differences between the two approaches for several taxonomic groups and seasons. As for spatial *β*-patterns, a proper comparison with the raw data could not even be done due to the (inevitably) limited number of eDNA sampling sites available. It is widely acknowledged that performing richness and diversity studies on raw eDNA data is problematic due to the large number of false negatives arising from the multiple steps of the metabarcoding procedure [Fukaya et al., 2022]. In eDITH, this aspect is accounted for via the assumption of geometric distribution of observed read number values for a given site and taxon. This allows detecting components of richness and diversity that would be otherwise overseen. Indeed, we found genus richness values estimated via eDITH to be often higher than those assessed from the raw data (Fig. 2).

Importantly, we found patterns of predicted *α*- and *β*-diversity to be strongly dependent both on the taxonomic group and season, which is in contrast with the predictions of Vannote et al. [1980] and Finn et al. [2011] of seasonally invariant and increasing *α*-diversity and decreasing *β*-diversity in the downstream direction, respectively. The only taxonomic groups for which patterns were found to be in accordance with these predictions are fish for *α*-diversity and bacteria for *β*-diversity. Overall, however, drainage area was found to be a poor (and often even non-significant) predictor of patterns of both diversity types. Indeed, occurrence of genera resulted to be correlated to a variety of environmental covariates, with considerable variation in importance and direction of the effect across seasons and taxonomic groups (see Fig. S2). On the same data set, Blackman et al. [2021a] assessed effect of drainage area and season on genus richness via mixed models (applied on the raw eDNA data), finding a positive, significant effect of drainage area for fish, and a significant effect for bacteria whose direction depended on the season. Our modelling approach came to a similar conclusion with respect to these groups, but additional negative, significant effects of drainage area for invertebrates in spring and summer were observed (Table 1).

Predicted patterns of invertebrates were found to be highly variant in both space and time, as spatial and temporal *β*-diversity values were the highest with respect to other taxonomic groups (Figs. 4, 5, Tables S1, S2). Indeed, most invertebrate genera were predicted to be located only in limited parts of the catchment and showed a marked temporal variability. Only 2.5% of the invertebrate genera (i.e., 2 out of 80, *Baetis* and *Eiseniella*) were predicted to be present in at least 25% of the same reaches across the three seasons (Fig. S5). In contrast, bacteria communities showed lower variability in both space and time, with a core group of genera that were found to be present in large portions of the catchment irrespective of the season: 9.6% of the detected genera (i.e. 27 out of 282) occupied at least 25% of the same reaches at all seasons (Fig. S5). Plausible explanations for limited variability in bacteria patterns are their limited dispersal abilities, and the fact that bacteria genera generally consist of a higher number of species than is the case for the other taxonomic groups, hence possible turnover at the species level is here hidden. Higher variability of invertebrate patterns as compared to bacteria is also supported by the fact that predicted invertebrate richness in spring explains less variation in richness in the other seasons as compared to bacteria (and partially to fish, see Fig. S4). Fish were found to be the most stable taxonomic group, with temporal *β*-diversity values much lower than those for invertebrates and bacteria (Fig. 5, Table S2), and spatial values lower than those for invertebrates and comparable to those for bacteria (Fig. 4, Table S1). This result is likely influenced by the limited number (12) of fish genera detected, and the ubiquity of the genus *Salmo*, which was predicted to be present in *>* 80% of the same reaches across the three seasons (Figs. S5, S6).

Overall, we see two main mutually non-exclusive explanations for these patterns and their limited overlap with previous models and findings [Vannote et al., 1980; Finn et al., 2011]. First, past predictions of riverine diversity patterns were often based on a small number of sites, often situated in a linear line-up along the main river stem. Thereby, these studies did not consider contributions of small-scale spatial dynamics from the dendritic river network, nor temporally fluctuating dynamics in organisms’ abundance and occurrence. Such time-invariant assumptions may be adequate for some groups, yet are likely not realistic for systems with a pronounced seasonality, including alternation between high and low flows or even desiccation, resulting in complex population dynamics. Second, our approach assumes an even detectability across space for the taxa considered. Likely, this strong assumption is at least to some degree violated, as eDNA-based data are known to be highly affected by stochasticity [Deagle et al., 2014; Elbrecht and Leese, 2015; Kelly et al., 2019] or may not be totally replicable, especially with generic primers as used here. Thus, some of the observed patterns may also reflect heterogeneity in the sampling procedure itself. While we cannot separate these processes, we are confident that the latter (uneven detectability) does not play a dominant role. Note also that possible temporal differences in detectability have been accounted for by the modeling procedure, since taxon distribution patterns were obtained separately for each season, and by using time-specific values for the hydrological variables. Therefore, despite eDNA shedding rates being dependent on environmental factors such as water temperature and metabolic activity [Jo et al., 2019; Thalinger et al., 2021] and hence arguably on season, these aspects do not affect our predictions.

In particular, seasonal differences in diversity patterns could be related to biological (when taxa actually change their abundance and/or spatial distribution across seasons) or methodological (when the likelihood to detect taxa changes across seasons) aspects. In the latter case, absence data may not be indicative of a true absence, but may be interpreted as an “ecological absence” (i.e. not ecologically relevant at that time point due to low density [Blackman et al., 2021a]). In our results, several observed patterns have a plausible biological explanation. First, the higher *α*-diversity of fish in low size reaches in summer as compared to the other seasons (Fig. 3) is possibly due to the migrating behaviour of several fish taxa during the spawning season, which, for widely found species in European temperate rivers, such as the gudgeon (*Gobio gobio*) and the common minnow (*Phoxinus csikii*), happens in late spring and early summer (see the predicted presence/absence maps for genera *Gobio* and *Phoxinus* in Fig. S6). Second, the lower *α*-diversity of invertebrates in autumn (Fig. 3) is likely related to the fact that many aquatic insects have already completed the aquatic part of their life cycle in autumn. Instead, the observed lower bacteria richness in summer compared to spring and autumn could have a methodological explanation: indeed, the total number of reads observed for bacteria in summer is intermediate with respect to those for spring and autumn, resulting however in a lower number of detected genera (198) with respect to spring (220) and autumn (214) (Fig. S1). This apparent mismatch between read number count and number of detected genera could be explained by the proliferation of some bacterial taxa in summer following temperature increase, which could mask the DNA from rarer taxa in the sequencing procedure, such that most (or all) amplification is biased towards the dominant taxa. Such patterns, similarly to primer bias, are well-known in metabarcoding studies [Kelly et al., 2019], and we acknowledge that the relatively low volume of water sampled may not have resulted in sufficiently saturated species accumulation curves.

The use of the eDITH model allowed an enhancement of our capability of interpreting eDNA data, enabling the extraction of patterns of *α*- and *β*-diversity at a much higher spatial resolution than what could be achieved by analyzing the eDNA data alone. Indeed, the spatial resolution at which model predictions are produced can be tuned freely, and this choice does not add any complexity to the model fitting procedure, as long as eDNA production rates *p*_*i*_ are expressed as a function of environmental covariates. In this application, we imposed a maximum reach length of 1 km, which resulted in a total of 1908 reaches, more than double than the number of reaches (760) used in a previous application in the same catchment [Carraro et al., 2020b]. However, it is important to note that a finer discretization of the river network would result in predicted richness values at the different reaches that would be more interdependent, which would require adequate tools (such as the ad-hoc statistical tests performed here) to analyze the resulting patterns. Another caveat in this respect, pointing at an opposite direction, is that too fine of a discretization might result in unrealistic small-scale differences in taxon patterns (e.g., presence-absence-presence predicted at a sequence of short, flow-connected reaches); a potential solution is offered by roughness-minimizing approaches borrowed by fluvial geochemistry in the analogous problem of predicting geochemical maps in catchments based on downstream water samples [Lipp et al., 2021].

In the present analysis, we used drainage area as the variable defining the spatial position of local communities (reaches) in the river network. Drainage area is the master variable controlling the bulk of hydrological and geomorphological features that shape a fluvial landscape (such as water discharge, velocity, river width and depth [Rodriguez-Iturbe and Rinaldo, 2001]) but also organic matter and nutrient availability [Bertuzzo et al., 2017; Helton et al., 2018; Jacquet et al., 2021], and hence also determine local habitat conditions. Thus, its use is, both from a hydrological as well as ecological perspective, highly suitable. Importantly, however, increasing *α*- and decreasing *β*-diversity patterns in river networks had previously been described mostly with respect to stream order [Vannote et al., 1980; Finn et al., 2011; Altermatt, 2013]; our choice of using drainage area is consistent with previous studies, as this variable varies predictably with stream order according to Horton’s law on drainage areas [Schumm, 1956]. It is likely, however, that our approach might have an impact on estimation of trends of spatial *β*-diversity. The partitioning of reaches into an “upstream” and a “downstream” group was roughly equivalent to comparing headwaters (i.e., reaches with stream order equal to 1) to all other reaches (Fig. S7). While pairwise Jaccard distances for reaches of stream order 4 or 5 could be lower with respect to those for more upstream reaches, it is unfeasible to statistically assess the magnitude of this trend for the different stream order values, because of the paucity of reaches with stream order ≥ 4 in the river network, and the fact that these tend to be connected by flow (which is likely to bias conclusions on *β*-diversity, see Methods). Note also that the predominance of headwaters with respect to reaches of high stream order is not limited to the catchment studied here, but is rather a universal feature of river networks [Horton, 1945].

We acknowledge that the herein assessed riverine biodiversity patterns do not cover sampling variability at the site level; it is indeed known that read number values vary substantially as a result of stochasticity in the sequencing process [Deagle et al., 2014; Deiner et al., 2015; Elbrecht and Leese, 2015], and thus our results contain some level of stochastic variation that we cannot control for. Higher robustness of eDITH predictions would be possible by incorporating true sampling replicates of the eDNA water samples, upon which the model fitting procedure is applied. Moreover, the predicted biodiversity patterns could be validated by comparing them with abundance estimates from direct observation of organisms. While this is feasible for fish (e.g. via electrofishing) and invertebrate (via kicknet sampling) groups, validation of bacterial patterns would be more complicated, as it would rely again on metabarcoding approaches from different supports (e.g. biofilm, although free floating bacteria communities can arguably not fully overlap with communities growing in benthic biofilm). In this respect, a possible improvement of the eDITH model would consist in the merging of eDNA and direct organismal observation data via joint-likelihood data integration methods [Miller et al., 2019]. Another conceivable expansion of the current analysis regards the use of taxon richness predictions provided by eDITH to foster food web analysis at an unprecedented spatial resolution (see Blackman et al. [2021a] for assessment of food web characteristic and functional diversity based on the raw eDNA data). This analysis seems particularly promising in the case study at hand, given the clear link that we observed between richness patterns of contiguous trophic level (Fig. S3). In conclusion, the eDITH approach, which transforms eDNA data into space-filling predictions of biodiversity, is generally applicable to any taxonomic group in riverine ecosystems and any temporal window, providing biodiversity predictions that can be used in the analysis of ecosystem processes, as well as to implement targeted (both spatially and temporally) conservation interventions. Predicted patterns of *α*- and *β*-diversity across taxonomic groups and seasons were not generally amenable to a simple increasing or decreasing trend in the downstream direction, but rather to a complex interplay of environmental variables, abiotic and biotic factors, highlighting the need for differentiated conservation approaches in riverine systems. A thorough assessment of biological communities in rivers cannot forego the integration of these aspects, and the eDITH model can be an adequate tool for such integrated analyses.

## 5 Acknowledgements

We thank Samuel Hürlemann, Xing Xing, Silvana Kaeser, Elvira Mächler, Roman Alther, Silvia Kobel and Aria Minder for help during field and laboratory work. Data were generated in collaboration with Jean-Claude Walser and the Genetic Diversity Centre (https://gdc.ethz.ch), ETH Zurich, Switzerland. Funding is from the Swiss National Science Foundation Grants No 31003A 173074 and PP00P3 179089 (to FA) and the University of Zurich Research Priority Programme on Global Change and Biodiversity (URPP GCB, to FA).

