## Supplemental Information for "Modelling eDNA transport in river networks reveals highly resolved spatio-temporal patterns of freshwater biodiversity"

#### Supplementary Tables

**Table S1:** Mean values of pairwise spatial $\beta$-diversity (calculated among all 1908*1907/2 pairs of sites). $\beta_{JAC}$: Jaccard distance; $\beta_{JTU}/\beta_{JAC}$: proportion of Jaccard distance explained by the turnover component. Partitioning of $\beta$-diversity is calculated via the *betapart* package [Baselga and Orme, 2012].

| | $\beta_{JAC}$ | | | $\beta_{JTU}/\beta_{JAC}$ | | |
|---|---|---|---|---|---|---|
| | spring | summer | autumn | spring | summer | autumn |
| Fish | 0.501 | 0.527 | 0.591 | 0.335 | 0.358 | 0.500 |
| Invertebrates | 0.752 | 0.772 | 0.764 | 0.823 | 0.853 | 0.803 |
| Bacteria | 0.499 | 0.635 | 0.560 | 0.753 | 0.805 | 0.711 |

**Table S2:** Values of temporal $\beta$-diversity averaged across the 1908 reaches. $\beta_{JAC}$: Jaccard distance; $\beta_{JTU}/\beta_{JAC}$: proportion of Jaccard distance explained by the turnover component. Partitioning of $\beta$-diversity is calculated via the *betapart* package [Baselga and Orme, 2012].

| | $\beta_{JAC}$ | | $\beta_{JTU}/\beta_{JAC}$ | |
|---|---|---|---|---|
| | spring-summer | spring-autumn | spring-summer | spring-autumn |
| Fish | 0.562 | 0.360 | 0.266 | 0.255 |
| Invertebrates | 0.754 | 0.742 | 0.866 | 0.777 |
| Bacteria | 0.694 | 0.659 | 0.831 | 0.909 |

### Supplementary Figures

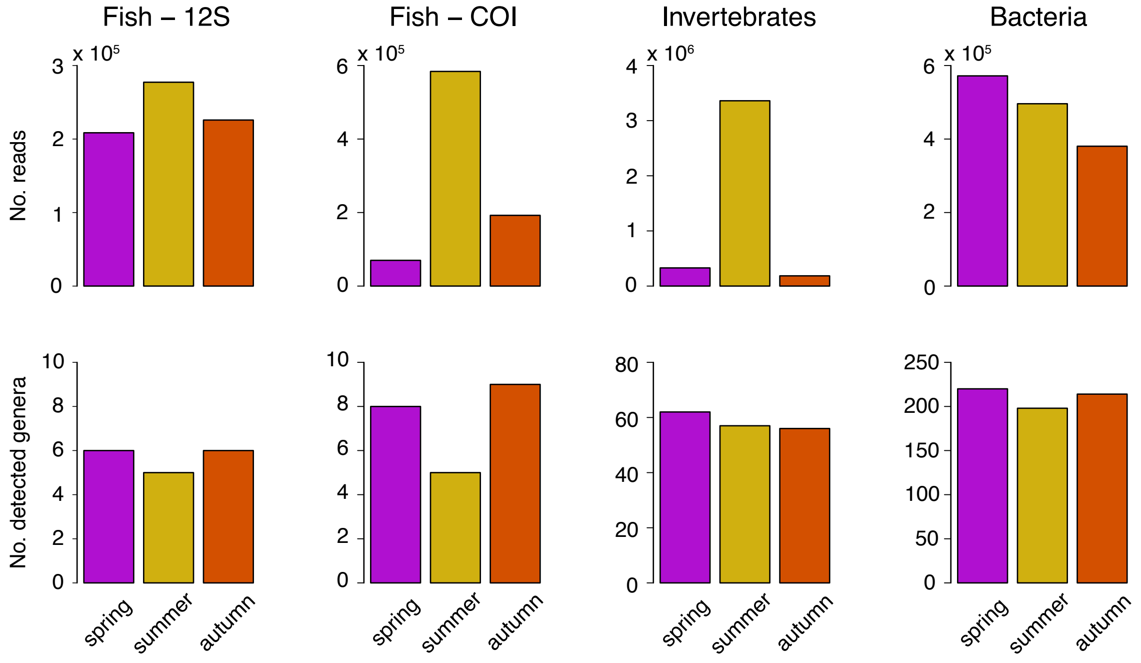


**Figure S1:** Summary of eDNA metabarcoding data pooled over sampling sites, and partitioned by taxonomic group, barcode region (for fish), and season. Note that here the data corresponding to fish genera *Barbus*, *Gobio* and *Phoxinus* are included in the histograms for both 12S and COI barcode regions.

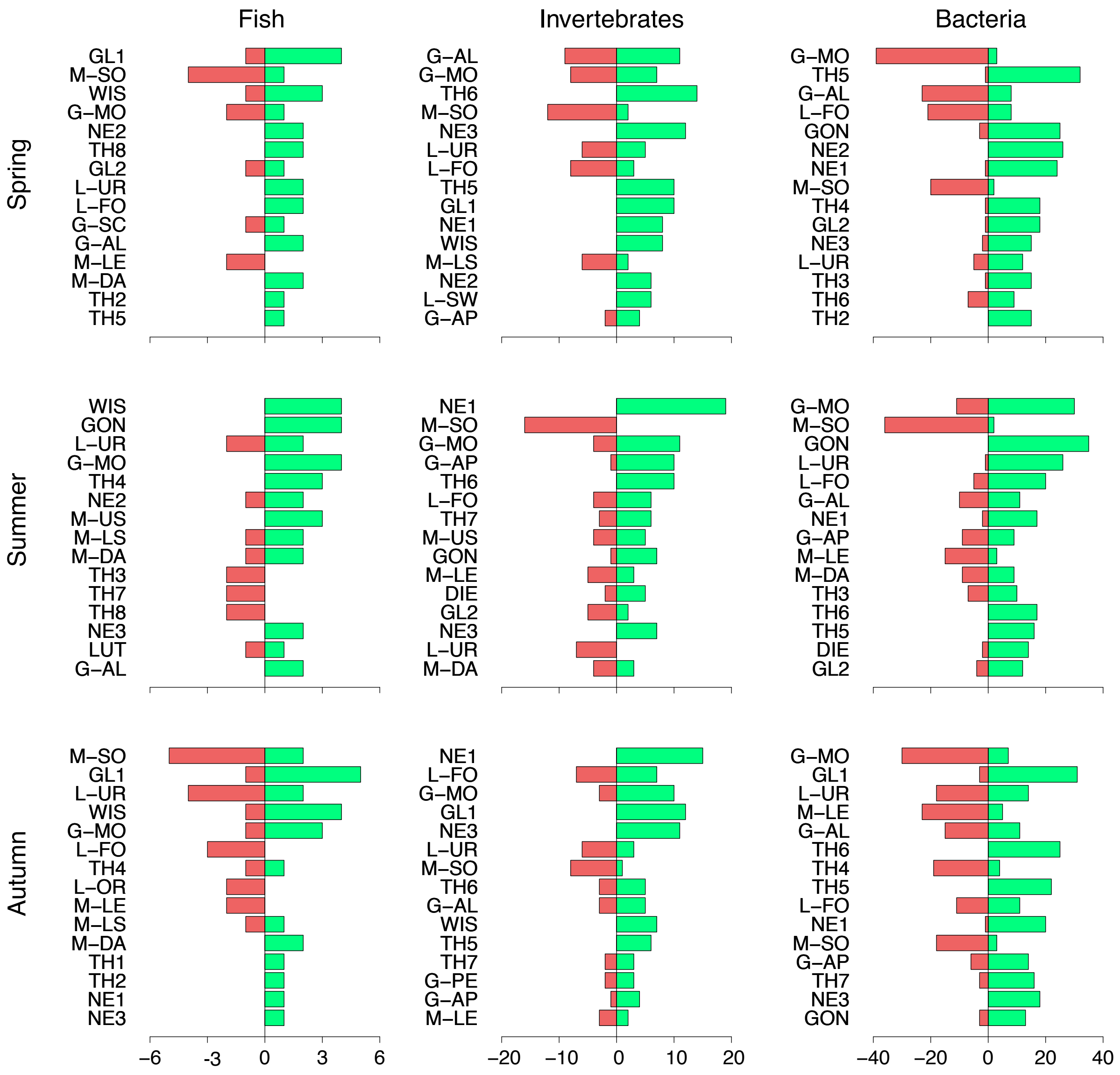


**Figure S2:** Significance of environmental covariates. The x-axis expresses the number of genera within a given taxonomic group and season for which a significant positive (green) or negative (red) effect was predicted by the eDITH model. Acronyms in the y-axis identify environmental covariates (see Supplementary Table 1 in Carraro et al. [2020] for the key). For graphical reasons, only the 15 (out of 35) most significant variables are shown. Positive (negative) significance is attributed if the 2.5$^{th}$-97.5$^{th}$ percentile range of the posterior distribution of the respective $\beta$ parameter was positive (negative). See Carraro et al. [2020] for details.

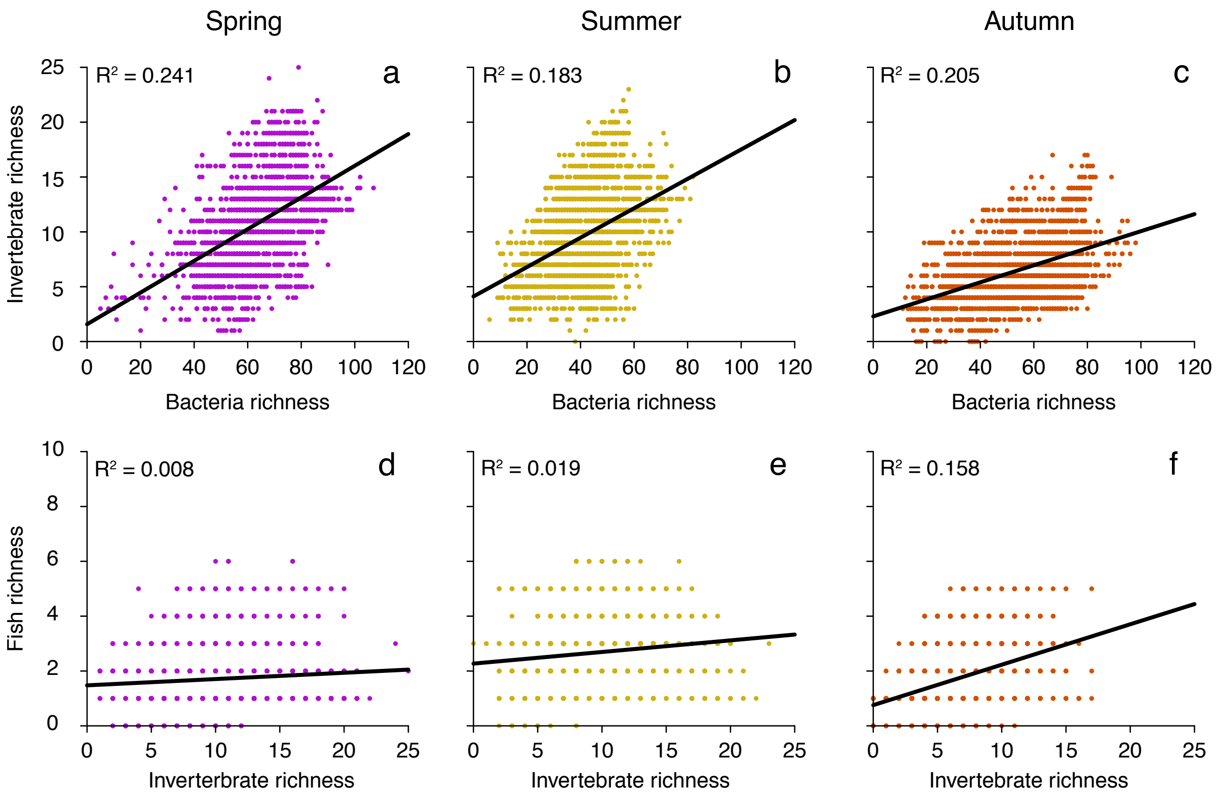


**Figure S3:** Relationship between patterns of $\alpha$-diversity with respect to different taxonomic groups (top row: bacteria richness vs. invertebrate richness; bottom row: invertebrate richness vs. fish richness). Black solid lines represent linear regressions. $R^2$ values are reported on the top-left corner.

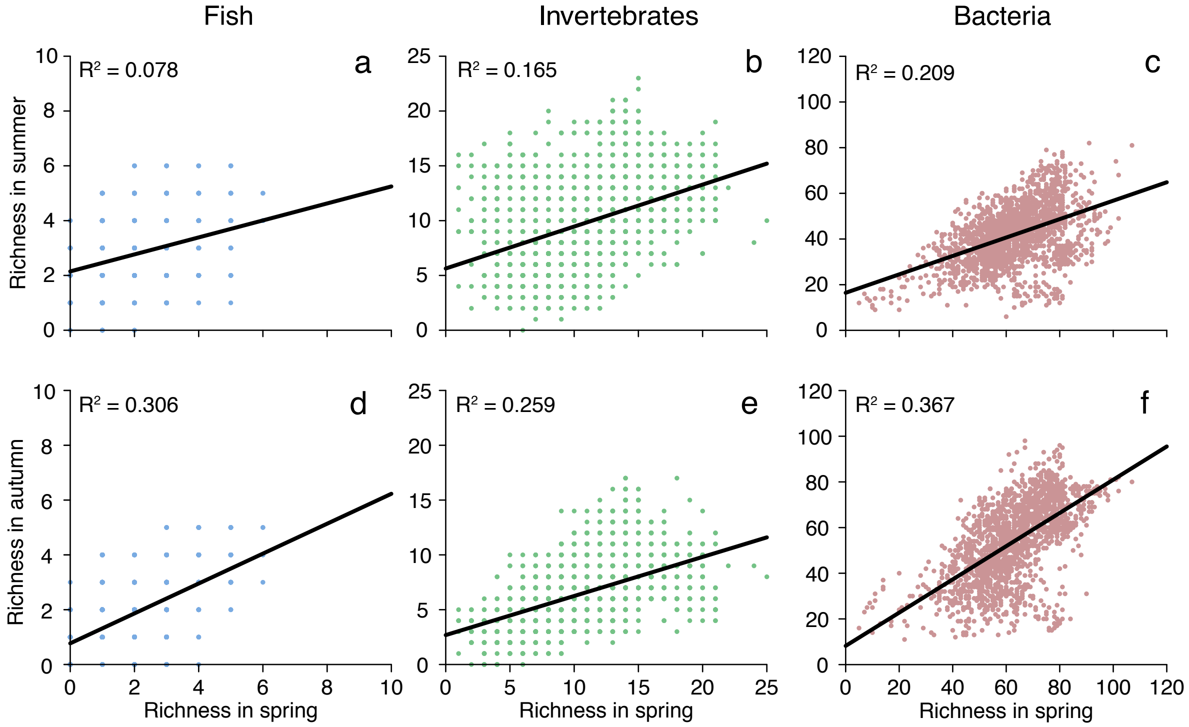


**Figure S4:** Relationship between patterns of $\alpha$-diversity with respect to different seasons (top row: richness in spring vs. richness in summer; bottom row: richness in spring vs. richness in autumn. Black solid lines represent linear regressions. $R^2$ values are reported on the top-left corner.

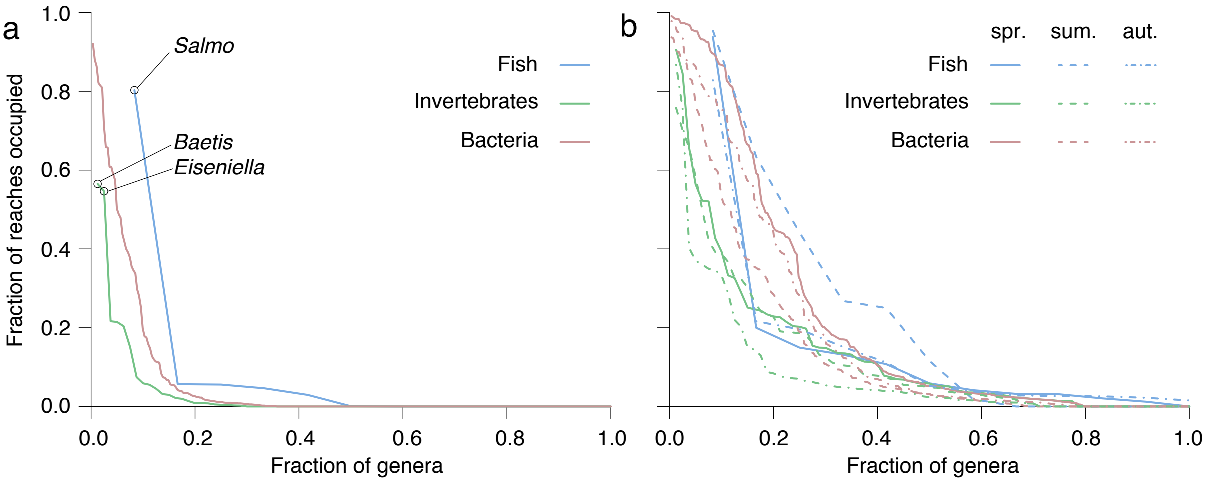


**Figure S5:** Distribution of genera for a given taxonomic group sorted by the fraction of reaches occupied as predicted by the eDITH model. a) Values pooled across seasons (here a taxon is considered present in a reach only if presence therein was predicted at all three seasons); b) values partitioned by season. Interpretation: a point $(x, y)$ on a curve expresses that there is a fraction $x$ of genera that are predicted to occupy a fraction $\geq y$ of reaches. In panel a, the most abundant fish and invertebrate genera are marked with dots.

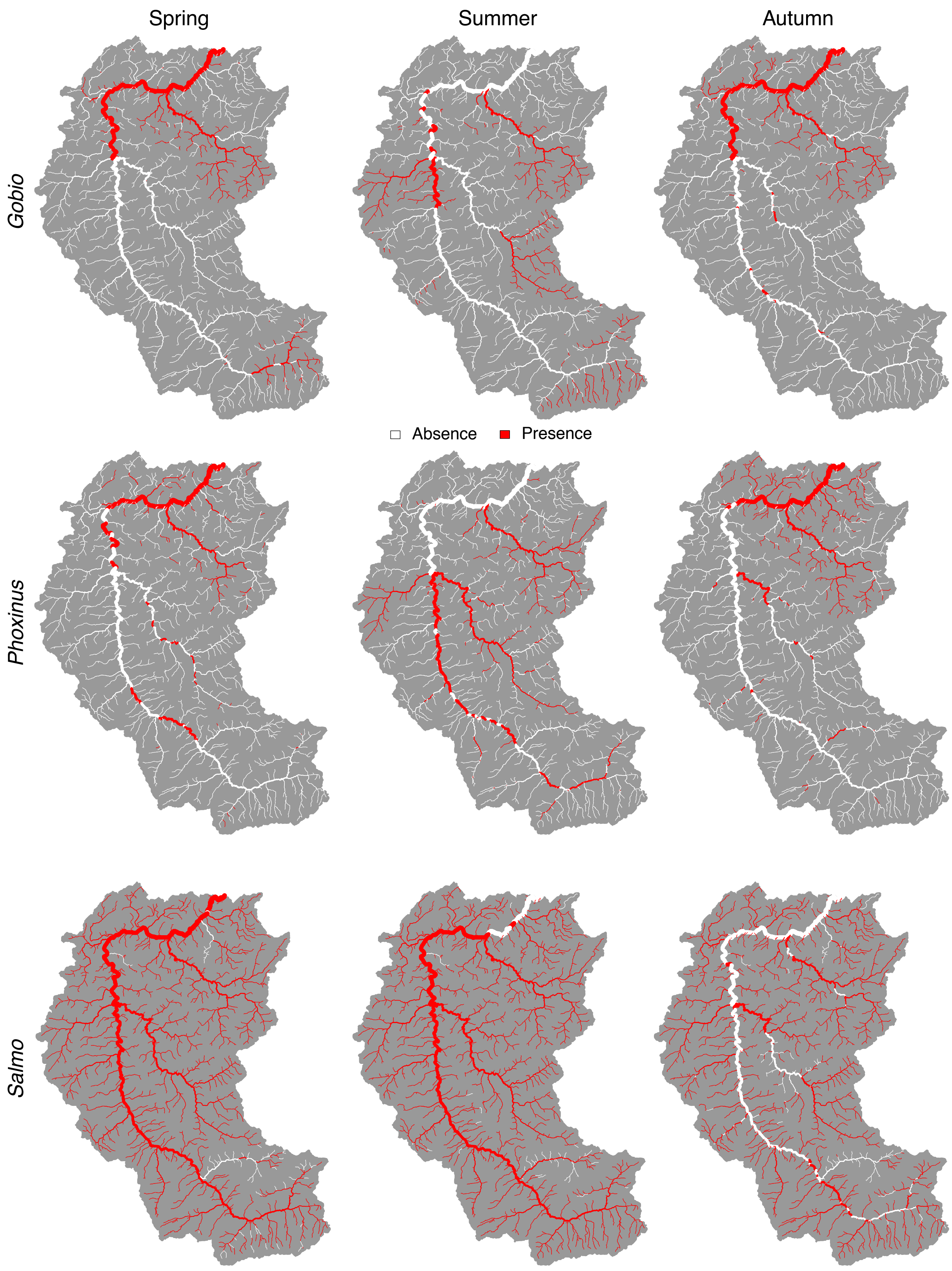


**Figure S6:** Presence-absence maps as predicted by the eDITH model for fish genera *Gobio*, *Phoxinus* and *Salmo* across the different seasons.

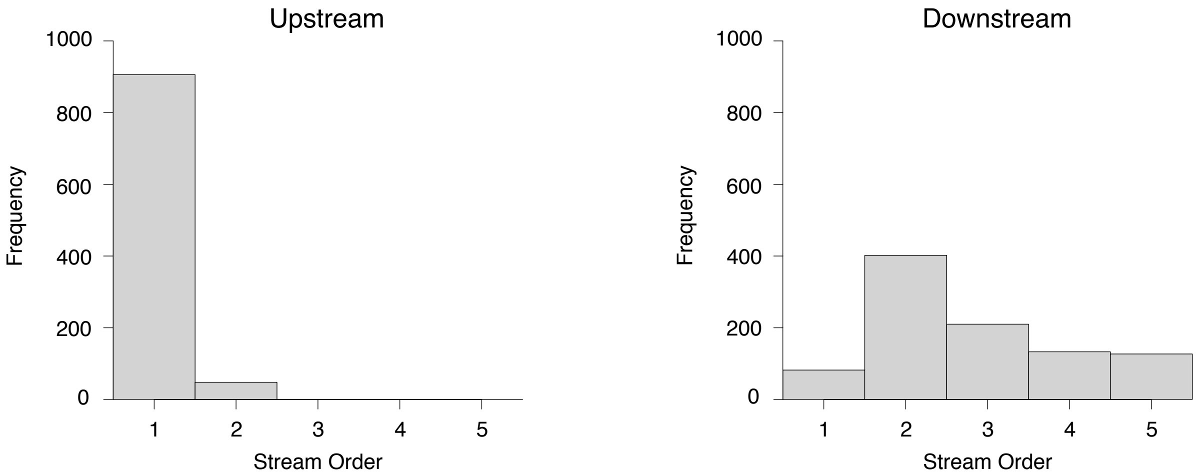


**Figure S7:** Distribution of stream order values within the groups used for analysis of spatial $\beta$-diversity patterns. "Upstream" ("Downstream") refers to reaches whose drainage area was lower (higher) than the median.
